# Characterization and mechanism of lead and zinc biosorption by growing *Verticillium insectorum* J3

**DOI:** 10.1101/404640

**Authors:** Chong-ling Feng, Jin Li, Xue Li, Ke-lin Li, Kun Luo, Xing-sheng Liao, Tao Liu

## Abstract

*Verticillium insectorum* J3 was isolated from a local lead-zinc deposit tailing, and its biosorption characteristics and reaction to the toxicities of different Pb(II) and Zn(II) concentrations were investigated. SEM, FTIR, a pH test and a desorption experiment were carried out to identify a possible mechanism. The biosorption of J3 presented an inhibition trend at low concentrations (25-75 mg L^-1^) and promotion at high concentrations (100-300 mg L^-1^). J3 absorbed Pb(II) prior to Zn(II) and produced alkaline substances, while mycelial and pellet morphology modifications were important for the removal of Pb(II) and Zn(II) under different stressful conditions (SEM results). Both intracellular accumulation and extracellular absorption may contribute to the removal of Pb(II) at lower concentrations (25-50 mg L^-1^), although mainly extracellular biosorption occurred at higher concentrations (75-300 mg L^-1^). However, Zn(II) bioaccumulation occurred at all concentrations assayed. *Verticillium insectorum* J3 may have evolved active defenses to alleviate the toxicity of heavy metals and proved to be a highly efficient biosorbent, especially for Pb(II) at high concentrations. This study is a useful reference for the development of biotreatment technologies to mitigate heavy metal waste.

## Introduction

With increased industrial and mining activities, contamination by heavy metals such as Pb and Zn seriously threatens both the environment and human health in Hunan. Lead (Pb) is a hazardous waste and is highly toxic to humans, animals, plants and microbes [1, 2]. Zinc is useful to humans at lower concentrations but is harmful at higher concentrations [3]. Bioremediation, characterized by lower costs and higher efficiency than physicochemical methods, has been an attractive option and is one of the hottest areas of research investigating the removal of heavy metals [4-6]. Many biomaterials, such as yeasts, bacteria, fungi and seaweeds, are capable of removing heavy metals or reducing their toxicity by directly or indirectly interacting with heavy metals[7, 8]. Some microorganisms are able to transform Pb(II) and Zn(II) through oxidation, reduction, methylation, and demethylation reactions [9]. Different functional groups on the biomass surface, such as carboxyl, hydroxyl, sulfhydryl and amino groups, also play important roles in the biosorption process [10].

Compared with dead microorganisms, metal-resistant living cells may demonstrate better removal of heavy metals due to continuous metabolic uptake after physical adsorption [11, 12]. Biosorption mechanisms are more complicated and change according to the type of heavy metal and cell viability. Living systems require special transport and handling mechanisms to keep them from contacting toxic metals in polluted environments [13-15]; many microorganisms reduce metal toxicity through bioprecipitation, and some have been shown to immobilize heavy metals by adsorbing metals on their cell surface or via intracellular accumulation [16, 17]. Naik *et al* [18] found that *Providencia alcalifaciens* bioprecipitates Pb(II) as lead phosphate and adsorbs metals onto its cell surface. Roane[19] isolated Pb-resistant *Bacillus megaterium* from metal-contaminated soils and found that this bacterium accumulated Pb intracellularly in the form of dense granules in the cytoplasm.

Mechanisms for heavy metal removal have been extensively studied. However, most researchers have focused on Pb(II) and Zn(II) resistance mechanisms for coping with high levels of heavy metal stress. These studies aimed to investigate the distribution of metal in cells, the role of extracellular compounds or the importance of intracellular sequestration, and they failed to fully take into account how microorganisms cope with toxic Pb and Zn during their growth and how defense systems respond to different initial metal concentrations. Moreover, the different roles of extracellular adsorption and intracellular accumulation, which are involved in the reduction of Pb(II) and Zn(II) toxicity, are still not clear. In the context of environmental stress and adaptation, it is of particular interest to understand how growing strains react to the toxicity of different Pb(II) and Zn(II) concentrations; such information would be useful for the development of fungi-based technologies to improve the degradation of metal pollution.

Strain J3 was isolated from a lead-zinc deposit tailing located in Zixing, Hunan, and identified as *Verticillium insectorum*. The strain survived and grew even in liquid medium with 500 mg L^-1^ Pb(II). The main objective of this study was to (a) investigate the biosorption characteristics of the strain under different conditions and assess Pb(II) and Zn(II) removal potential; (b) discuss the surface interactions between the growing strain and Pb(II) and Zn(II) under different stressful conditions; and (c) explore the roles of intra- and extracellular sequestration at lower or higher Pb(II) and Zn(II) concentrations.

## Materials and methods

### Microorganism, medium and culture conditions

The locally tolerant bacterium J3 was isolated from a lead-zinc mine tailing in Hunan, China, and was tested in this study. Spore suspensions of J3 were typically inoculated into modified Martin’s liquid (glucose 10.0 g, peptone 5.0 g, MgSO4·7H2O 0.5 g, water 1000 mL, pH 4.0±0.1, MGL) medium, and shake-flask culturing was performed with 120 rpm agitation at 30 °C. Then, lead and zinc at different densities were added according to the requirements for individual experiments.

### Chemicals

Nitrate salts (Pb(NO_3_)_2_, Zn(NO_3_)_2_·6H_2_O) of analytical grade were used to prepare 5000 mg L^-1^ stock metal ion solutions, which were diluted to create metal ion solutions for adsorption. Before the microorganisms were mixed, the pH value of each test metal solution was adjusted to the desired value with 1 mol L^-1^ NaOH or 1 mol L^-1^ HNO_3_.

### Biosorption experiments during growth

Experiments investigating the effects of pH, heavy metal concentrations, time and biosorbent dose on biosorption in growing J3 cells were conducted. The solution pH ranged from 3.0-8.0 in 1 mL of inoculum, and the reaction mixtures were shaken on an orbital shaker at 120 rpm and at 30 °C for 8 d. Similarly, the inoculum volume (0.2-2 mL), initial metal ion concentration (0-300 mg L^-1^), and contact time (3-8 d) were investigated. In the biosorption experiments, unless otherwise stated, the initial metal concentration, pH, temperature, and inoculum volume were 100 mg L^-1^, 4.0, 30 °C and 1 mL, respectively. Growing cells were harvested by filtration and rinsed with ddH_2_O three times after culture, and then vacuum freeze-drying was conducted to obtain the dry biomass.

The concentrations of Pb(II) and Zn(II) in the filtrate were determined by atomic absorption spectrophotometry (AAS) after filtration. The values of the biosorptive capacity and removal ratio of Pb(II) and Zn(II) were calculated from the following equations:

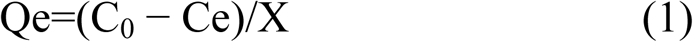

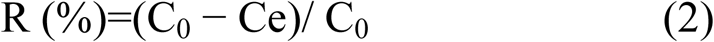

where Qe is the equilibrium Pb(II)/Zn(II) concentration of the biosorbent (mg g^-1^ dry cell); R is the removal ratio of Pb(II)/Zn(II); C_0_ and Ce are the initial and residual metal concentration (mg L^-1^), respectively; and X is the biomass concentration (g dry cell L^-1^).

### Scanning electron microscopy (SEM)

To investigate the effects of Pb(II) and Zn(II) on the morphology of mycelium pellets and hyphae, SEM analysis was employed. Metal-free (control) and metal-loaded (100-300 mg L^-1^) biosorbents were vacuum freeze-dried for 24 h. The pellets of metal-free and metal-loaded biosorbents were opened with a sterile knife, and the interior and external features were observed, while hyphae were ground down into flour for observation. Pre-treated samples were coated with Au via vapor deposition prior to observation by SEM (Hitachi S4100, Tokyo, Japan).

### FT-IR analysis

Infrared spectra of metal-free (control) and metal-loaded (50-300 mg L^-1^) biosorbents were obtained using a 330 spectrometer (Nicolet Avatar, USA). Dried and crushed samples mixed with KBr were analyzed with a spectrophotometer in the range of 4000-400 cm^-1^. The background was subtracted from the sample spectra before analysis.

### pH test after culture with various concentrations of heavy metals

The effects of metal ion concentration on biosorption were studied as described in section 2.3. The initial pH was adjusted to 4.0, and the pH of the liquid medium was tested by pH meter after the cells were harvested.

### Desorption experiments

Living J3 cells were cultured in different metal iron concentrations (0-300 mg L^-1^) as described in section 2.3. Then, the dry biomass was desorbed with 1 M HNO_3_ solution; after 12 h of gentle agitation, the biomass was harvested by centrifugation (8000 rpm, 10 min). The metal concentration in the supernatant was determined by AAS. The desorption efficiency, D (%), was calculated according to Eq (3).

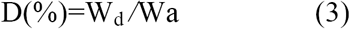

where W_d_ is the amount of Cd desorbed (mg), and Wa is amount of Cd adsorbed (mg).

## Results and discussion

### Influencing factors

#### Effects of initial pH

Each type of microorganism has a particular pH range for growth, so the initial pH of the growth solution plays a significant role in the biosorption process[20], affecting the activity of functional groups (phosphate, carboxylate and amino groups) and the chemical properties of metals (Fig 1). Low pH also leads to the competition of metal ions for binding sites [21].

**Fig 1.**
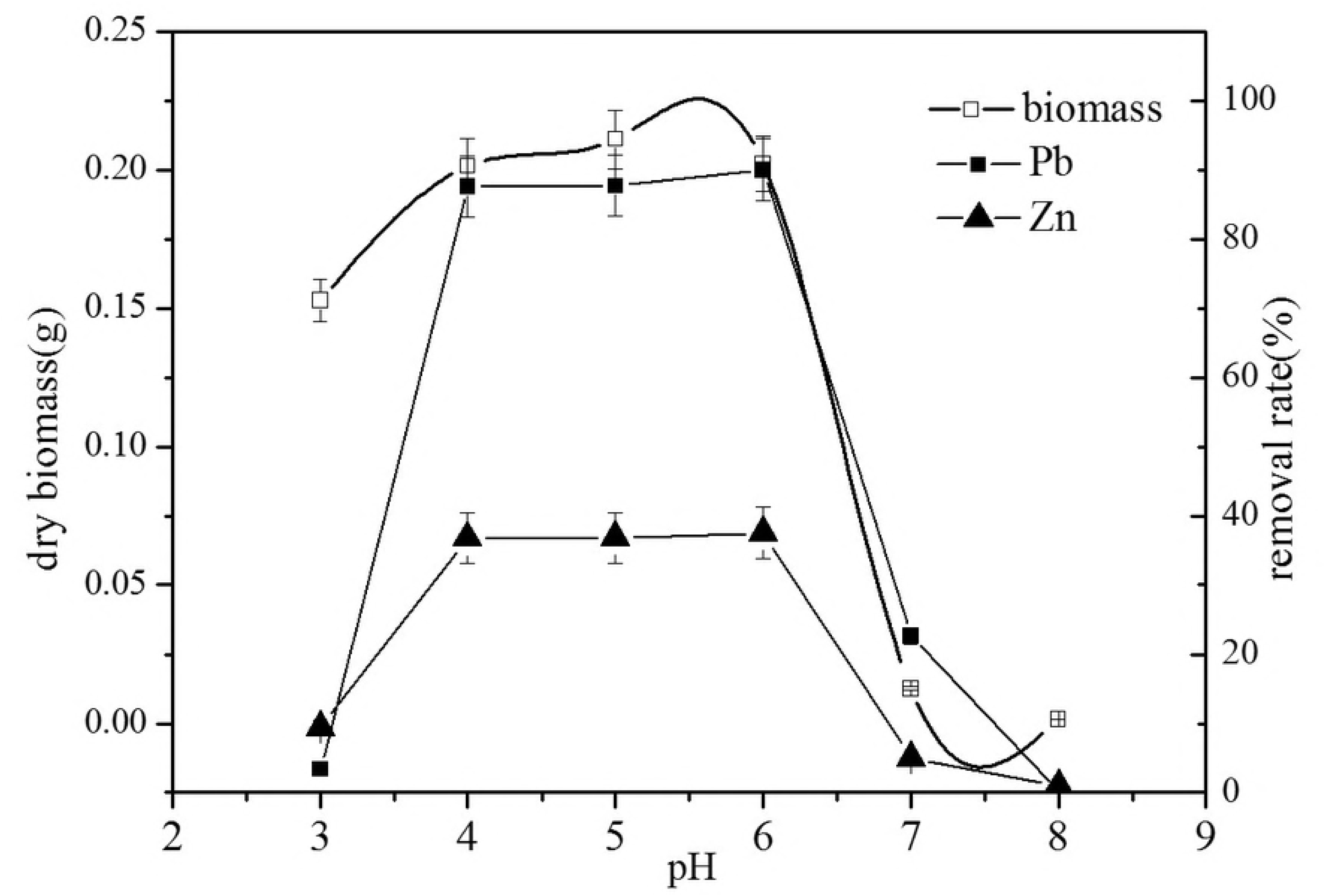
The effects of initial pH on biosorption.

As shown in Fig 1, the biosorption capacity of the two metal ions was much lower at low pH (3.0), although the biomass of J3 showed no significant difference at pH 4.0, 5.0 and 6.0. Lower biosorption yield is attributed to the large quantity of hydrogen ions competing with metal ions for available binding sites [22]. At pH values of 4.0 to 6.0, the removal rates of Pb(II) and Zn(II) were approximately 90% and 36%, respectively. This result may be attributable not only to the increased biomass of J3 but also to the increased number of binding sites on the cells at pH values above the isoelectric point, which is beneficial for the removal of Pb(II) and Zn(II) [23]. At pH 7.0 and 8.0, the biomass of J3 was lower and accompanied by a rapid decrease in the biosorption capacities of Pb(II) and Zn(II). Although the maximum biosorption yield (90.02%) of J3 was identified at an initial pH of 6.0, a similar biosorption yield and faster growth of J3 were obtained at an initial pH of 4.0. Therefore, subsequent experiments were carried out at pH 4.0.

#### Effects of initial concentration

The maximum biomass was obtained at initial Pb(II) and Zn(II) concentrations of 25 mg L^-1^, which indicated that the growth of J3 was promoted at low concentrations of metal ions. However, the removal rate of Pb(II) was merely 17.6% under these conditions (Fig 2). The removal rate of Pb(II) sharply increased with increasing concentrations from 25 mg L^-1^ to 100 mg L^-1^, demonstrating that biosorption was not determined by the biomass but was related to the concentrations of Pb(II) and Zn(II). The biomass was maintained at approximately 0.17 g, and the removal rate of Pb(II) was approximately 90% when the initial Pb(II) and Zn(II) concentrations ranged from 75-250 mg L^-1^. Although the biomass decreased at 300 mg L^-1^, the removal rate remained steady. This result might be attributed to the enhancement of fungal metabolism to maintain energy in response to Pb stress., Fungi might have evolved active defense mechanisms to alleviate the toxicity of metals[10, 24]. However, the mechanism was not completely clear and required further investigation; thus, subsequent experiments were carried in this study.

**Fig 2.**
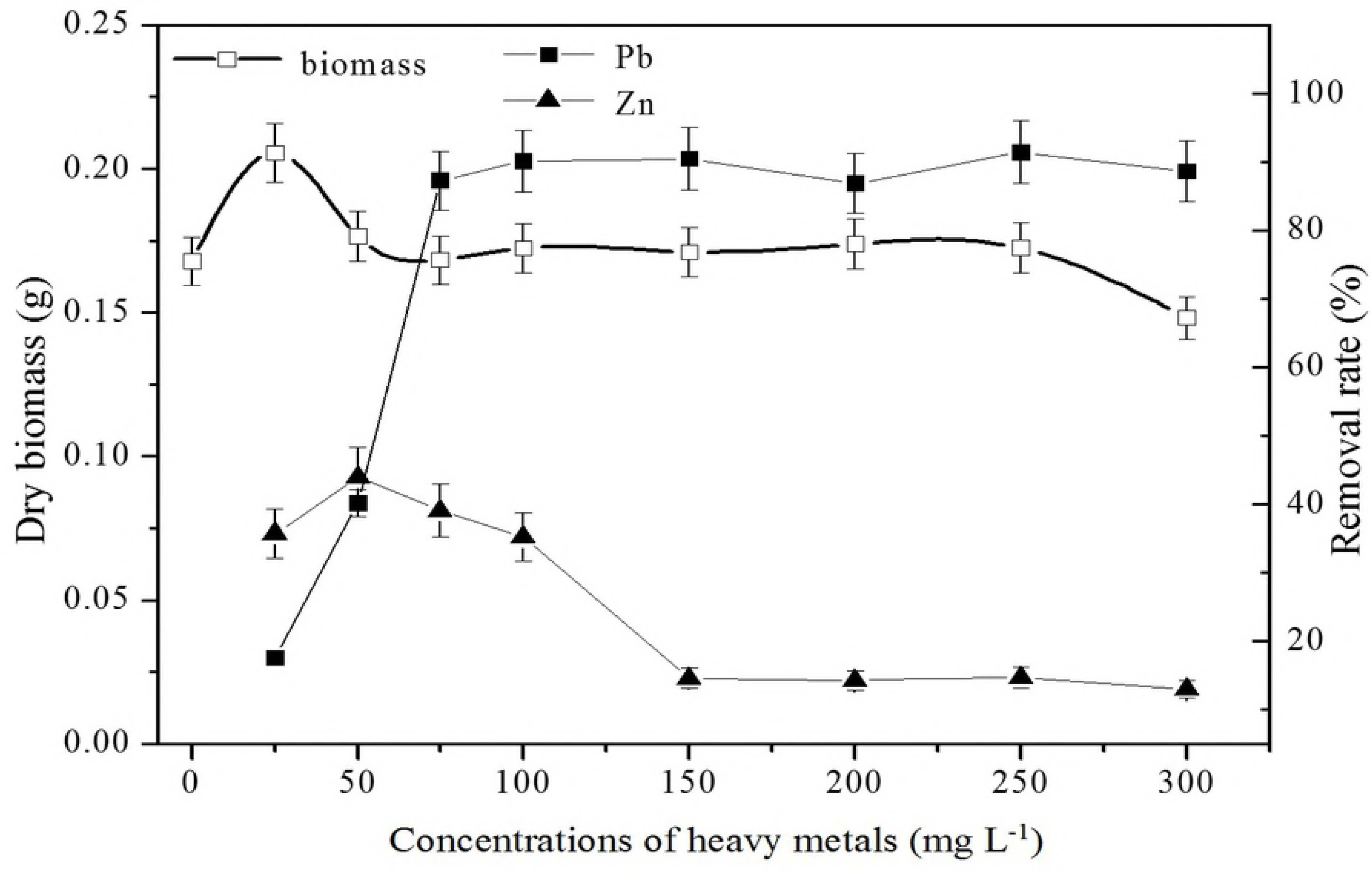
The effects of the initial concentrations of Pb(II) and Zn(II) on biosorption.

Zinc is an essential nutrient and has no toxic effect on growth at low concentrations. The removal rate increased from 35.7% to 43.9% as the initial concentration increased from 25 mg L^-1^ to 50 mg L^-1^. After that, the removal rate decreased until it remained steady at approximately 15%. The removal rate of Pb(II) was much higher than that of Zn(II), which might be due to competition for adsorption[25, 26].

#### Effects of inoculum dose

The amount of inoculum is also an important parameter for biosorption. There was a significant difference between the absorption of Pb(II) and Zn(II). The removal rate of Pb(II) was as high as approximately 90%, and this value remained stable when the inoculum dose was higher than 0.5 mL. However, the removal rate of Zn(II) increased coordinately with increasing inoculum dose (Fig 3). Two possible scenarios were hypothesized based on this result: (1) J3 absorbed Pb(II) prior to Zn(II), in other words, the absorption of Zn(II) happened after a certain level of Pb(II) saturation was achieved; or (2) the increased removal rate of Zn(II) might be due to the increased inoculum dose, meaning J3 needs more Zn(II) because it is an essential element for growth.

**Fig 3.**
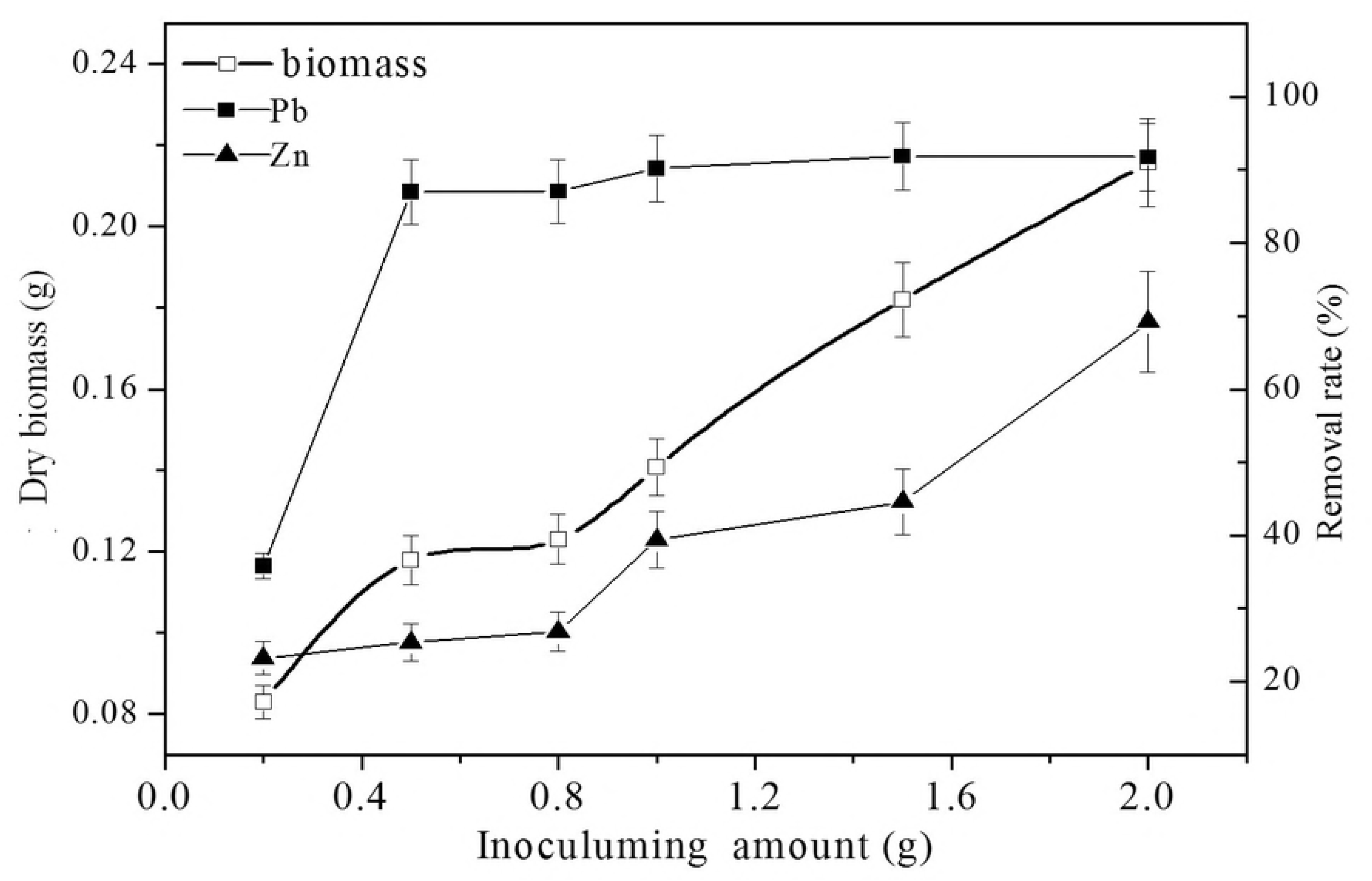
The effects of inoculum amount on biosorption.

#### Effects of absorption time

The effects of absorption time on Pb(II) and Zn(II) were also determined and are shown in Fig 4. It was noted that the Pb(II) and Zn(II) absorption process could be divided into three stages. Absorption was slow due to a lag phase in J3 growth at early time points. Subsequently, as more biomass accumulated (logarithmic growth phase), the removal rate increased rapidly. Finally, it reached adsorption equilibrium on the 8^th^ day, and approximately 90% Pb(II) was removed, while the Zn(II) removal rate was 33%. However, the maximum removal rate for Zn(II) occurred on the 7^th^ day. The results indicated that absorption mainly happened during the logarithmic growth phase, and the absorption of Pb(II) and Zn(II) was competitive, such that the absorption of Pb(II) was prioritized over that of Zn(II). This result agreed with the results described in sections 3.2 and 3.3.

**Fig 4.**
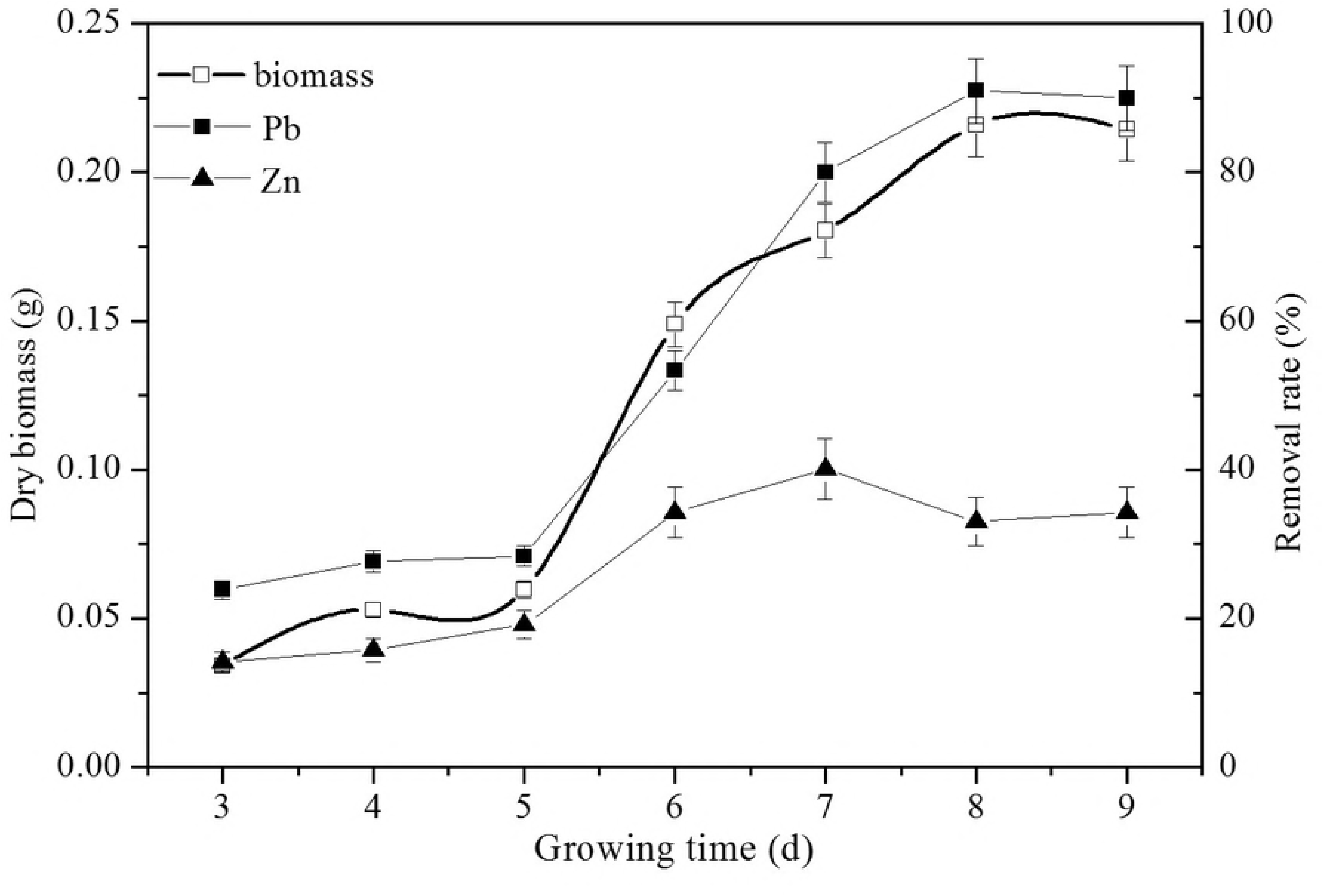
The effects of growth time on biosorption.

#### SEM analysis

SEM micrographs of growing cells before and after the biosorption of Pb(II) and Zn(II) are shown in Fig 5. Compared with the control (a), SEM showed that the surfaces of the pellets (b) were covered by a mass of gelatinous substance; the reason for this might be the secretion of extracellular polymeric substances or metabolic products of J3 complexes with Pb(II) and Zn(II) through cell-surface binding, ion exchange or precipitation[16]. As shown in panels b to d, the surfaces of metal-loaded hyphae looked vague and distorted and seemed to be damaged by increasing levels of heavy metal ions. Hypha morphology changed gradually. At lower Pb(II) and Zn(II) concentrations, the hyphae were smooth and resisted biosorption. However, at higher Pb(II) and Zn(II) concentrations, the cells warped and aggregated, which could improve extracellular adsorption capacity, thereby raising tolerance to lead and zinc. This morphological change may be important for Pb(II) and Zn(II) removal, as similar responses induced by heavy metals have been found by other researchers[27, 28].

**Fig 5.**
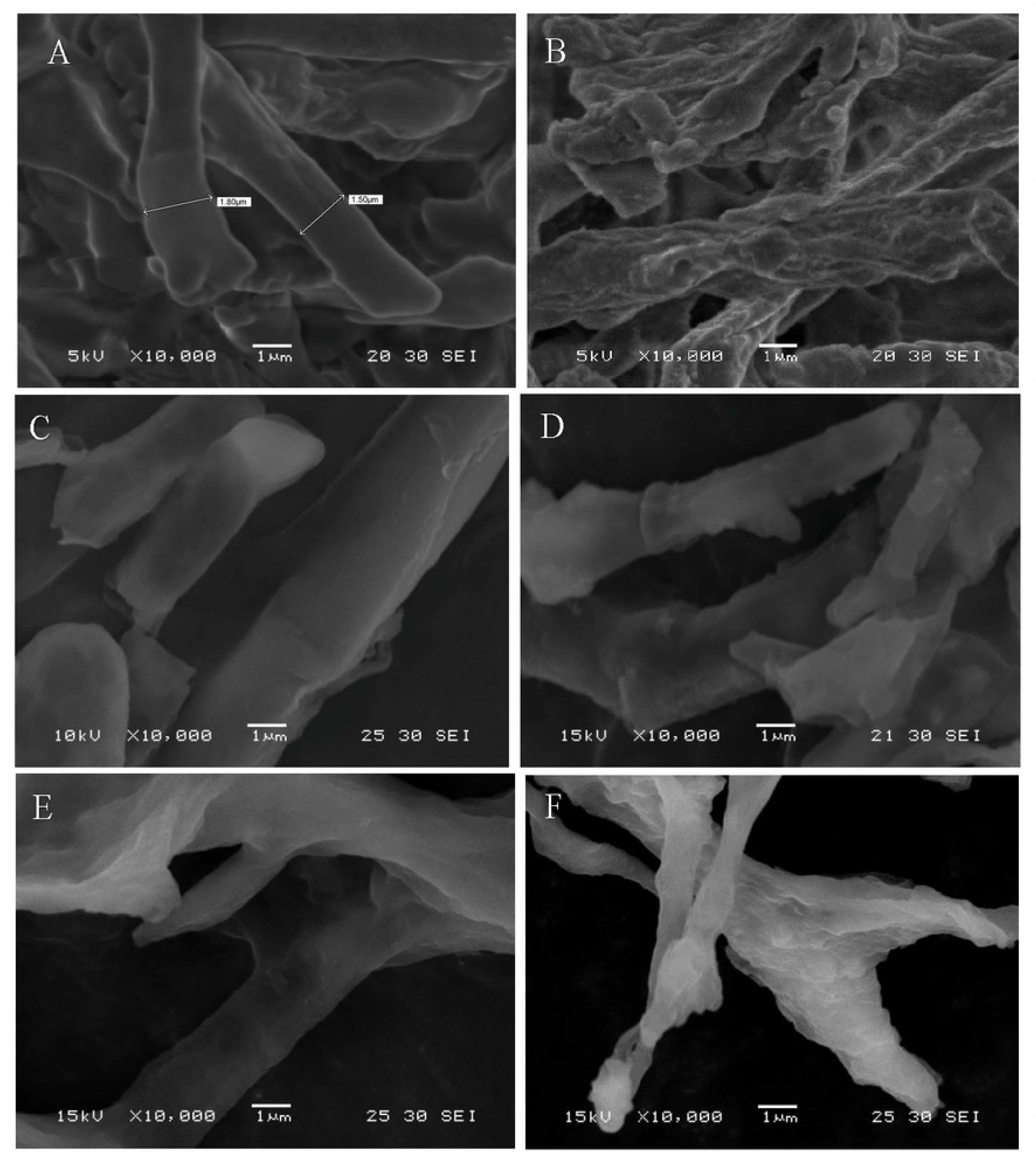
SEM micrograph of growing *Verticillium insectorum* J3 under different Pb(II) and Zn(II) concentrations. (a) Metal-free pellet (control); (b) metal-loaded pellet (300 mg/L); (c) metal-free hypha (control); (d), (e), (f) metal-loaded hyphae (100, 200, 300 mg L^-1^, respectively)

Additionally, the surfaces of pellets from Pb(II)- and Zn(II)-free liquid media were sparse, smooth and loose, whereas densely entangled hyphae were found after exposure to Pb(II) and Zn(II). This coincided with a finding in an earlier study regarding the mycelial morphology of *Phanerochaete chrysosporium*. Baldrian[29] reported that the addition of a heavy metal (Cd) led to the formation of dense hyphae caused by an increase in the lateral number per branch point and a decrease in the distance between branch points. In this study, pellets without Pb(II) and Zn(II) were solid with a diameter of approximately 1 mm, but pellets with Pb(II) and Zn(II) were hollow with a diameter of approximately 3 mm. This result indicated that heavy metals affected pellet formation by J3. It is unclear whether this phenomenon represents a survival mechanism for J3 or is the result of poisoning. Further studies will be needed.

#### FTIR spectral analysis

To confirm the functional groups involved in the biosorption process, FTIR analysis was carried out. The absorbance spectra for the control cells and those loaded with Pb(II) and Zn(II) are shown in Fig 6 (A-E); a significant shift in the adsorption peaks can be observed.

**Fig 6.**
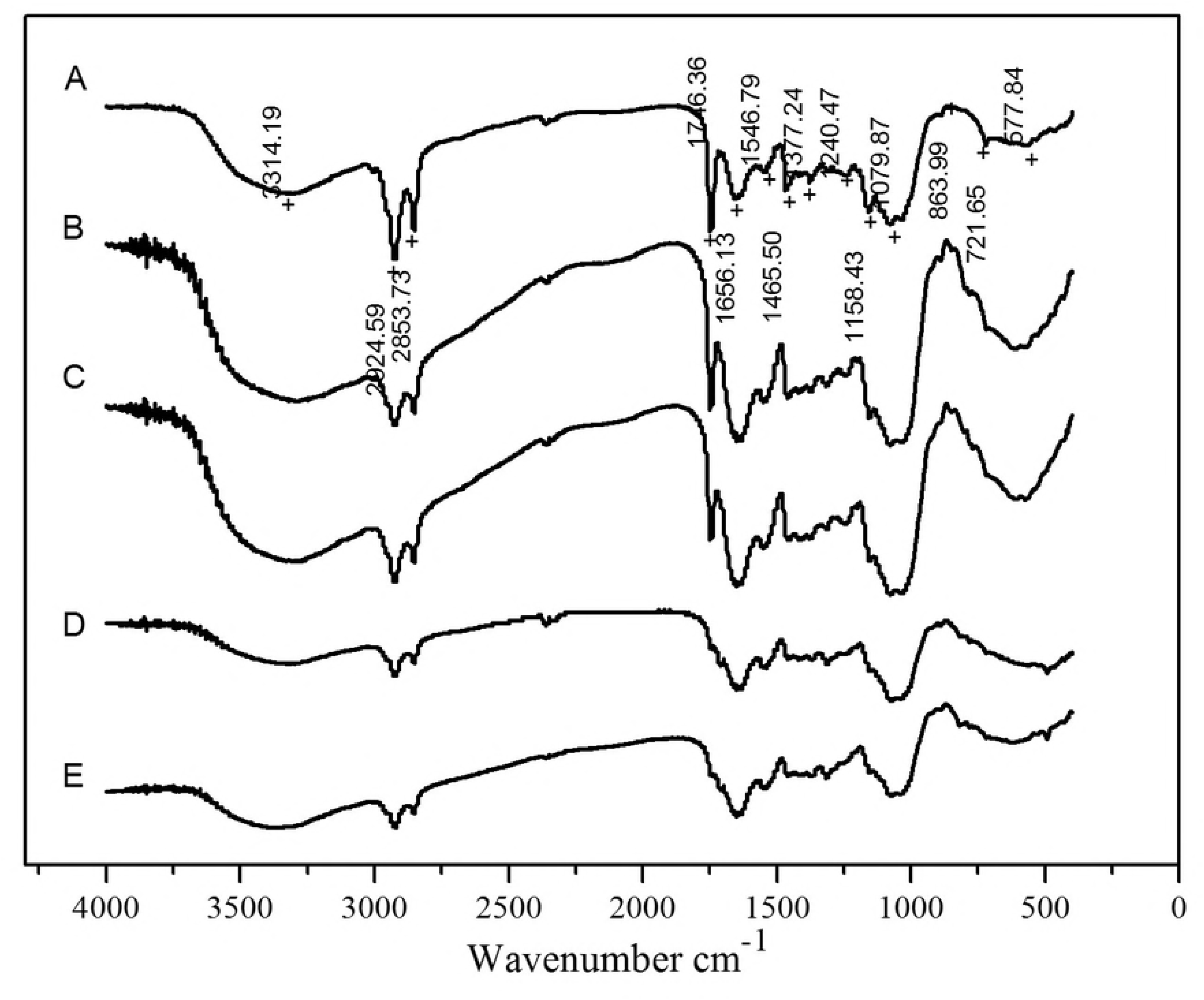
FTIR spectra of growing *Verticillium insectorum* J3 cells under different Pb(II) and Zn(II) concentrations. A is the control treatment; B, C, D, E are metal-loaded J3 cells (50, 100, 200, 300 mg L^-1^, respectively)

At the peak at approximately 3314.19 cm^-1^ (A), the FTIR spectra of J3 cells loaded with Pb(II) and Zn(II) (B, C, D, E) indicated band shifts to 3285.99 cm^-1^, 3301.58 cm^-1^, 3307.86 cm^-1^ and 3371.31 cm^-1^. These bands were attributed to the stretching vibration of O–H and N–H groups in polysaccharides, proteins and fatty acids. The peaks of B and C were significantly increased comparing with the control group, while the adsorption peaks of D and E were reduced and wider, which may be due to the large numbers of complex hydroxyl groups that coordinated with lead and zinc ions in the D and E groups.

The peaks at approximately 2924.59 cm^-1^ and 2853.73 cm^-1^ gradually diminished with increasing metal concentrations, and the bands shifted slightly, which was attributed to the stretching vibration of C-H groups from polysaccharides and proteins. The peak observed at 1746.36 cm^-1^ (A) was assigned to amide I found in proteins and peptides; potentially due to the stretching vibration of C=O groups, this peak was weaker in B and C and disappeared in D and E. In addition, the absorption band at 1656.13 cm^-1^ (A) was assigned to typical amides I and II and exhibited small shifts after the adsorption of heavy metal ions. The peak strengthened in B and C and then weakened in D and E. The other peak observed at 1546.79 cm^-1^ (A) was assigned to amide II and was attributed to -NH bending vibration or -CN stretching vibration. The peak shifted to 1548.5 cm^-1^, 1538.61 cm^-1^, 1538.46 cm^-1^, and 1538.46 cm^-1^ in B, C, D, E, respectively. According to the above analysis, the absorption peaks of -CH, amide I and II demonstrated significant shifts after interactions with Pb(II) and Zn(II), and polysaccharides, fats and the amide groups of proteins may play important roles in this process.

In particular, the peaks at 1240.47 cm^-1^ (Ar-O stretching) and 1158.43 (C-O stretching) shifted in groups B and C and disappeared in D and E. However, the peak observed at 1079.87 cm^-1^ (C-O stretching), which was assigned to aliphatic ether, was obviously stronger in B and C and weaker in D and E, although it was still stronger than the control group. As J3 was identified as *Verticillium insectorum*, this suggested that ethers not only participate in the adsorption of Pb(II) and Zn(II) but also may be related to pest resistance, although further studies are needed.

The above observations revealed that the -NH, -CH, -OH, C=O and C-O groups might participate in the Pb(II) and Zn(II) biosorption process. The results showed that the actions of many groups on fungal cell walls varied by concentration. Compared with control group A, the peaks in B and C exhibited variable regulation, as did those in D and E. The adsorption intensities of many groups increased in B and C relative to the control group and then decreased in D and E, although they were still stronger than A. At the same time, some other groups disappeared in D and E, potentially because metabolism or metabolic products were changed at various concentrations of Pb(II) and Zn(II).

#### pH test after culture at various heavy metal concentrations

In this experiment, the initial pH was 4; however, the final pH was >4.4 at the end of culture, with or without the presence of heavy metals (Fig 7). Compared with the control group, the pH increased obviously at concentrations ranging from 25-100 mg L^-1^. The maximum value (5.16) was observed at 50 mg L^-1^. Then, the pH value decreased with increased heavy metal concentrations. Borremans *et al* [30] found that the Pb-resistant bacterium *Ralstonia metallidurans* removed Pb from solution by increasing the pH to 9 during bacterial growth. Changes in pH indicated that J3 can produce alkaline substances, and iron exchange may occur at higher concentrations. Some microbes can precipitate metals through the production of organic bases. This ability might represent a method to complex Pb(II) and Zn(II) as insoluble precipitates, as observed by scanning electron microscopy (SEM), thus alleviating the toxicity of heavy metals. This result also agreed with the variations in O-H stretching revealed by FTIR spectra analysis.

**Fig 7.**
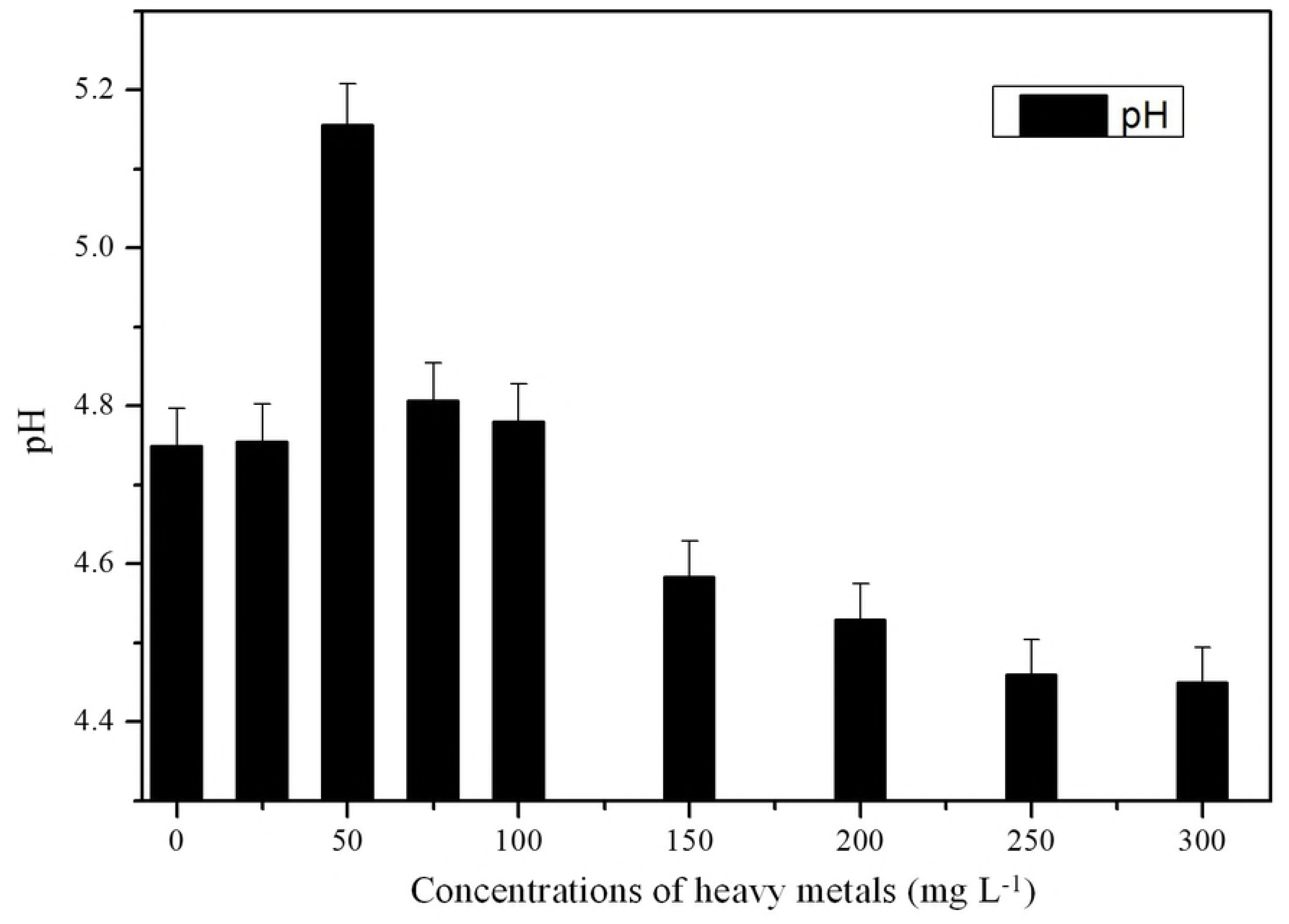
Changes in pH after biosorption at different concentrations.

#### Desorption experiments

In theory, the adsorption of heavy metal ions by a microbe surface is reversible. The ions can be desorbed from the microbe cell surface at low pH. In this study, 1 M HNO_3_ was selected as the desorption agent. The desorption of Pb(II) and Zn(II) from heavy-metal-loaded biomass is shown in Fig 8.

**Fig 8.**
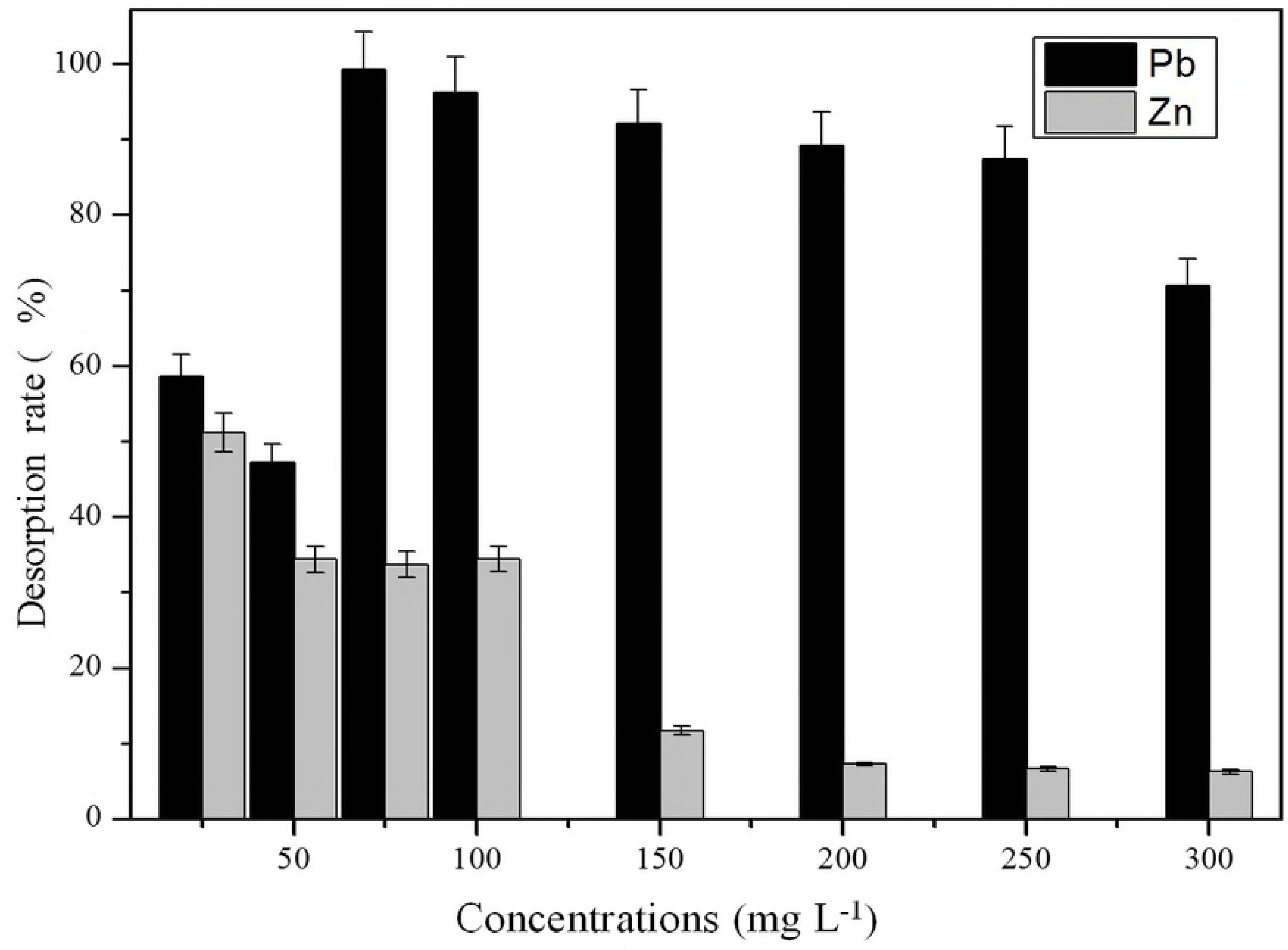
Desorption efficiency at different heavy metal concentrations.

The desorption rate of Pb(II) was 58.5% and 47.2% at Pb(II) concentrations of 25 mg L^-1^ and 50 mg L^-1^, respectively, and the desorption rate of Pb(II) was maintained above 70% at concentrations ranging from 75 mg L^-1^ to 300 mg L^-1^. This result suggested that both intracellular accumulation and surface absorption play important roles at low concentrations, while surface absorption was the major removal mechanism at higher concentrations (75-300 mg L^-1^). This is similar to the results reported by Huang *et al*[31]. However, the desorption rate of Pb(II) decreased after it reached maximal values at 75 mg L^-1^, according to the results described in section 3.2, and the removal rate remained stable. This result indicated that as more Pb(II) gets into the cell, J3 biomass decreases, as observed at 300 mg L^-1^ (Fig 2).

The desorption rate of Zn(II) was below 50% at all times and decreased with increasing concentration. This result suggested the bioaccumulation plays a leading role in the biosorption process. Gao *et al* [32] reported that a moderate dose of zinc could help antagonize the toxicity of lead in immature rats, but more studies are needed to determine whether the intracellular accumulation of Zn(II) results in decreased lead toxicity in <u>microorganism</u>s.

## Conclusions

The growth of J3 under Pb(II) and Zn(II) stress showed a promotion trend at low concentrations (25 mg L^-1^) and inhibition at high concentrations (300 mg L^-1^). An absorption experiment showed the absorption of Pb(II) prior to Zn(II). J3 absorbed Pb(II) and Zn(II) through cell surface binding, intracellular accumulation or precipitation (SEM results). The FTIR results suggested that -NH, -CH, -OH, C=O and C-O groups participate in the Pb(II) and Zn(II) biosorption process, and J3 produced alkaline substances. The morphological changes in pellets and hypha, functional groups, and pH suggested that J3 evolved active defense mechanisms to alleviate the toxicity of heavy metals. Both intracellular accumulation and extracellular absorption may play important roles in the removal of Pb(II) at lower concentrations (25-50 mg L^-1^), although mainly extracellular biosorption occurred at higher concentrations (75-300 mg L^-1^). However, bioaccumulation may play a leading role in the biosorption process of Zn(II). Overall, *Verticillium insectorum* J3 altered the biosorption mechanism under lower or higher Pb(II) and Zn(II) concentrations and proved to be a highly efficient biosorbent, especially for Pb(II). These results provide a useful reference for the development of biotreatment technologies for heavy metal-polluted waste.

## Acknowledgments

The authors thank Yi Zhang, Ph.D., and Wei-zhen Wang, Ph.D., for help with the experiments.

